# Predicting Kinase-Specific Phosphorylation Sites with Pretrained Protein Language Models

**DOI:** 10.1101/2025.03.12.642740

**Authors:** Mahdi Pourmirzaei, Farzaneh Esmaili, Kai Chen, Mohammadreza Pourmirzaei, Mohsen Rezaei, Duolin Wang, Dong Xu

## Abstract

Accurately predicting kinase-specific phosphorylation sites remains difficult due to the diversity of kinases and the context-dependent nature of substrate recognition. Importantly, aberrant kinase overactivation is a hallmark of many cancers including colorectal, gastric, liver, and breast tumors where dysregulated kinase signaling promotes malignant transformation, tumor progression, and therapy resistance. This underscores the clinical importance of understanding kinase-substrate relationships and precisely mapping phosphorylation events. In this paper, we introduce two complementary sequence-based architectures that operate directly on full-length substrate and kinase sequences. Stage 1 extends a task-agnostic prediction method, named Prot2Token, to jointly support three tasks: kinase-group classification from substrate sequences alone, kinase-substrate interaction prediction, and kinase-specific phosphorylation-site prediction while incorporating a self-supervised decoder pretraining task that predicts amino-acid positions from encoder embeddings. This pretraining substantially strengthens site prediction. Stage 2 specializes the architecture for phosphorylation-site prediction by replacing causal decoding of Prot2Token with a bidirectional one, yielding further gains. On standard benchmarks, the specialized model consistently outperforms widely used baselines. Beyond in-distribution evaluation, across both in-distribution and zero-shot settings of understudied dark kinases, we show the sign of zero-shot kinase-specific phosphorylation-site prediction capability. Together, these results indicate that jointly modeling substrate and kinase sequences provides a straight-forward, scalable approach to state-of-the-art, zero-shot-capable phosphorylation-site prediction.

## 1 Introduction

Post-translational modifications (PTMs) are crucial processes that regulate cellular functions and biological processes [1]. Among these modifications, protein phosphorylation is one of the most significant. It involves the attachment of a phosphate group to specific amino acids *serine* (`S`), *threonine* (T), or *tyrosine* (`Y`) rendering the target protein phosphorylated [2]. This process is crucial for signal transduction, cellular communication, and the regulation of various biological activities. Dysregulation of phosphorylation leads to severe diseases, including cancer, neurodegenerative disorders, and Alzheimer’s disease [3, 4]. Kinases, a group of specialized enzymes, catalyze this process by transferring phosphate groups to proteins [5, 4], and kinases in cancer are significantly more likely to have phosphorylation site (p-site) mutations compared to controls. However, the prediction of kinase-specific phosphorylation^‡^ is challenging due to the vast diversity and structural complexity of kinases, as well as the dynamic nature of protein interactions within cells. Each kinase recognizes specific sequence motifs, but variations in these motifs, context-dependent factors, and lack of abundant high-quality datasets make accurate predictions difficult [6].

In recent years, numerous approaches have been developed to predict kinase-specific p-sites, leveraging computational techniques such as machine learning and deep learning [7]. However, most of the existing methods share significant limitations. Many rely on ensemble approaches that use several, or in some cases hundreds, of models to achieve acceptable accuracy [8, 9]. While this may improve predictive performance, it introduces inefficiencies in computational resource usage and scalability. Additionally, these methods typically focus on peptides with lengths ranging from 7 to 15 amino acids, which represent only a small fraction of possible protein sequences [10–12]. This narrow focus not only fails to cover the full range of protein sequences but also risks losing critical structural information, which is essential for understanding kinase-substrate interactions at a more comprehensive level. As a result, the predictive performance of these methods lacks generalization across diverse protein contexts, limiting their applicability in real-world biological scenarios.

To address these limitations, we reformulate kinase-specific phosphorylation as three interconnected tasks: (i) *kinase group classification, kinase-substrate interaction prediction*, and *kinase-substrate psite prediction*.

In the first stage, inspired by *Prot2Token* [13], we cast all three tasks as next-token prediction problems and develop a unified framework that couples an autoregressive transformer decoder with the ESM-2 [14] pre-trained protein language model (PLM). This design enables direct end-to-end learning from substrate and kinase sequences (Figure 1). We further introduce a self-supervised pretraining phase to initialize the decoder weights, which substantially boosts performance in p-site prediction compared to the randomly initialized *Prot2Token* architecture.

**Figure 1.**
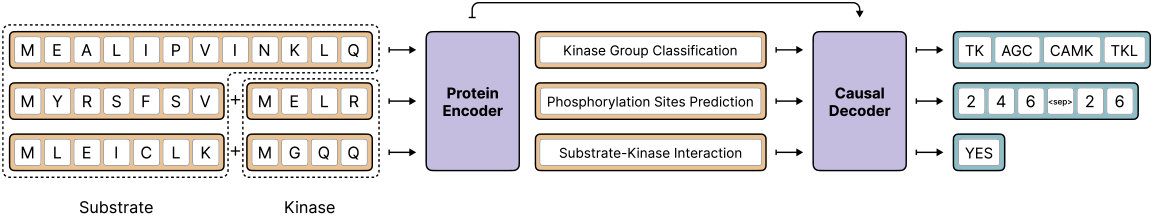
Overview of our framework for kinase-related predictions from substrate and/or kinase sequences. The model supports three tasks: *(i) kinase group classification* from substrate sequences, *(ii) kinase-substrate interaction prediction*, and *(iii) kinase-substrate p-site prediction*. A protein encoder produces sequence embeddings, which are fed into a causal decoder for task-specific outputs. Due to data leakage concerns, each task is trained independently; the diagram illustrates a unified view for conceptual clarity.

In the second stage, we derive a streamlined variant of this architecture specialized for kinase-specific p-site prediction only (Figure 3). This specialized model not only achieves a substantial performance gain over state-of-the-art methods but also exhibits the first evidence of *zero-shot* generalization—predicting p-sites for entirely unseen kinase sequences.

The contributions of this paper can be summarized as follows:

1. We extend the *Prot2Token* framework to jointly address three kinase-substrate related tasks: *kinase group classification* from substrate sequences alone, formulated as a multi-label learning problem; (ii) *kinase-substrate interaction prediction* using paired substrate and kinase sequences, enhanced by hard negative sampling inspired by data pruning [15] and contrastive learning [16]; and (iii) *kinase-substrate p-site prediction* using a self-supervised pre-training phase to initialize the autoregressive decoder, leading to notable gains in identifying kinase-specific p-sites.
2. We design a streamlined architecture specialized for kinase-substrate p-site prediction, achieving state-of-the-art results. We further evaluate this model on both in-distribution and out-of-distribution understudied kinases, providing evidence of zero-shot capability on entirely unseen kinase sequences during training.

Throughout this paper, we use the terms substrate sequence and protein sequence interchangeably, treating them as equivalent.

### 1.1 Related Work

The prediction of kinase-specific p-sites has been explored using various machine learning and deep learning approaches to advance our understanding of PTMs and their regulatory roles in cellular processes [11, 9, 17]. Computational models leverage sequence-based features, physicochemical properties, structural characteristics, and evolutionary information [7]. In contrast, deep learning methods aim for end-to-end learning by directly capturing complex patterns from raw data [18, 17]. Several computational models address p-site prediction, with some focusing broadly on PTM detection [19–21], and others dedicated solely to kinase-specific p-sites [11, 9, 22, 7]. Broadly, existing research in this domain falls into two main categories: kinase-specific p-site prediction and protein-kinase interaction prediction. The former focuses on identifying p-sites for specific kinases, while the latter aims to determine interactions between kinases and their substrate proteins. A detailed discussion of these categories and related work is provided in Appendix A.1.

## 2 Method

### 2.1 Stage 1

The architecture of this stage is based on *Prot2Token* where a causal (autoregressive) transformer decoder *T*_***ψ***_, whose parameters are indicated by ***ψ***, is connecting to a pre-trained bidirectional PLM, referred to as the encoder *G*_***θ***_, whose parameters are denoted by ***θ***. The substrate sequences *s* with an optional kinase sequences *k* are concatenated together and encoded by the *G*_***θ***_, producing residue embeddings *G*_***θ***_(*s and k*) ∈ ℝ^*M* ×*d*^. The decoder *T*_***ψ***_ integrates PLM representation through crossattention and a task token prompt *E*, after which, a linear binary classification head *C*_***ϕ***_, whose parameters are shown by ***ϕ***, produces residue level logits to perform next token classification on vocab sizes (Equation 1).

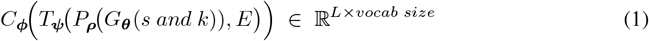

To control the distinct prediction of each task, we use separate tokenizers and embedding tables for the encoder and decoder (Figure 5). The autoregressive transformer factorizes the joint probability of a sequence *x* = (*x*_1_, *x*_2_, …, *x*_*m*_) into a product of conditional probabilities. Training proceeds by minimizing the negative log-likelihood of the observed tokens, where *θ* denotes the model parameters. Employing a causal mask, each token *x*_*m*_ can only attend to the tokens *x*_1_, …, *x*_*m*− 1_, thereby enforcing the autoregressive property and allowing the model to learn contextual representations of the preceding sequence.

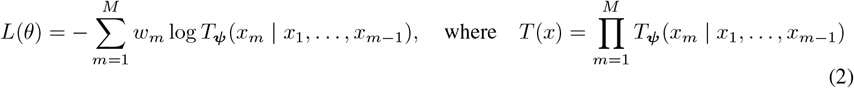

*Prot2Token* extends this standard autoregressive objective by introducing token-level weights *w*_*m*_ to regulate each token’s influence on the loss. Specifically, we set *w*_1_ = 0 so that predicting the first token (the prompt) does not affect the loss, while for *m* ≥ 2, *w*_*m*_ can be adjusted, granting non-prompt tokens varying degrees of importance. Concretely, the revised training objective demonstrated in Equation 2, where each weight *w*_*m*_∈ [0, ∞) is a user-defined parameter for the token *x*_*m*_. This setup enables flexible fine-tuning by emphasizing specific tokens of interest while removing the prompt token from the loss computation (by assigning it zero weight). Further details on the architecture are provided in Appendix A.2.

#### Tokenization of labels

We follow the tokenization framework introduced in *Prot2Token* to convert target labels into discrete tokens. Specifically, we use a non-hierarchical multi-label classification scheme for kinase group prediction, binary classification for protein-kinase interaction, and the same approach from the original work for both protein-kinase p-site and self-supervised tasks.

### 2.1.1 Self-Supervised Pre-Training

Our initial experiments in p-site prediction revealed that directly fine-tuning the *Prot2Token* model yielded suboptimal accuracy, even when testing various label formatting strategies. We hypothesized that this limitation stemmed from the randomly initialized decoder, which lacks the inductive biases required to interpret protein sequence information. As shown in Figure 2, the distribution of p-sites is sparse and highly imbalanced, making it difficult for an untrained decoder to learn meaningful positional representations from such a weak signal. This underscored the need to instill these priors through a dedicated self-supervised pre-training phase.

**Figure 2.**
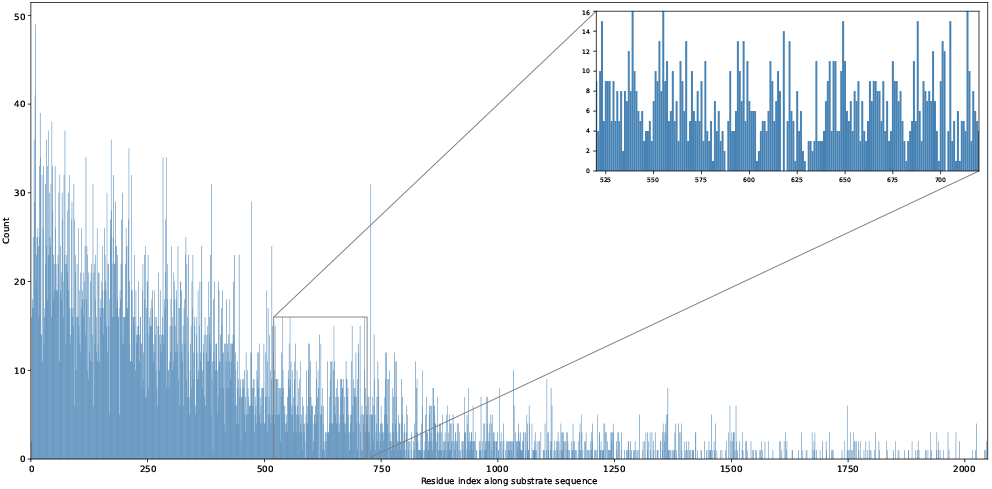
Distribution of p-site indices in the training set (n = 11 449 sites drawn from 5 694 sub-strates). Bars correspond to single-residue positions and cover indices less than 2 049; a further 176 extremely rare sites at higher indices are omitted for clarity. The histogram is highly imbalanced: a handful of low-index positions dominate, while most indices receive only a few or zero examples. This long-tail sparsity means the decoder’s embedding table sees too little signal at many positions to learn meaningful representations, particularly in the higher-index region where runs of zero counts are common.

To address this, we introduced a self-supervised pre-training strategy for the decoder, aiming to instill biologically meaningful priors before downstream fine-tuning. The core idea is to train the decoder to predict the positions of specific amino acids within given the representation of the protein encoder, thus enabling the model to learn position-aware residue representations without the need for manual annotations. For example, given a protein sequence such as `MSGLSNYT`, the task would be to identify the indices of each occurrence of a specified amino acid (e.g., `S`: positions 2, 5). We designed twenty such tasks, each corresponding to one of the twenty standard amino acids.

Self-supervised samples were generated automatically, making the approach scalable and costeffective. In particular, incorporating auxiliary prediction tasks for a broader set of amino acids, beyond the canonical phosphorylation targets (`S, T`), potentially provides a generality for different downstream site prediction type tasks.

### 2.2 Stage 2

This stage introduces a specialized p-site prediction architecture, which utilizes the *Stage 1* architecture and incorporates positional inductive biases for each amino acid by replacing the causal attention of the decoder with a bi-directional one and employing a shared-weights PLM to encode kinase and substrate sequences separately, and avoiding the decoder pre-training stage. Technically, substrate sequences *s* and kinase sequences *k* are tokenized and passed through the same pre-trained PLM *G*_***θ***_. Yielding an embedding *G*_***θ***_(*s or k*) *∈* R^*M* ×*d*^ whose dimensionality *d* generally differs from the decoder dimension *d*.

To align these feature spaces we introduce a learnable *linear projector P*_***ρ***_ : R^*M* ×*c*^ *→*R^*M* ×*d*^, parameterized by ***ρ***, producing *P*_***ρ***_ *G*_***θ***_(*s or k*) *∈* R^*M* ×*d*^ for the cross-attention layers of the same decoder *T*_***ψ***_. The fused representation is then passed to the binary classifier head *C*_***ϕ***_. Because *G*_***θ***_ is pre-trained on large protein sequence corpora, the architecture (Equation 3) can technically predict binding sites for entirely new kinase-substrate combinations, including those that have not been encountered during training (Figure 3).

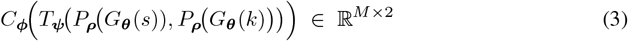

**Figure 3.**
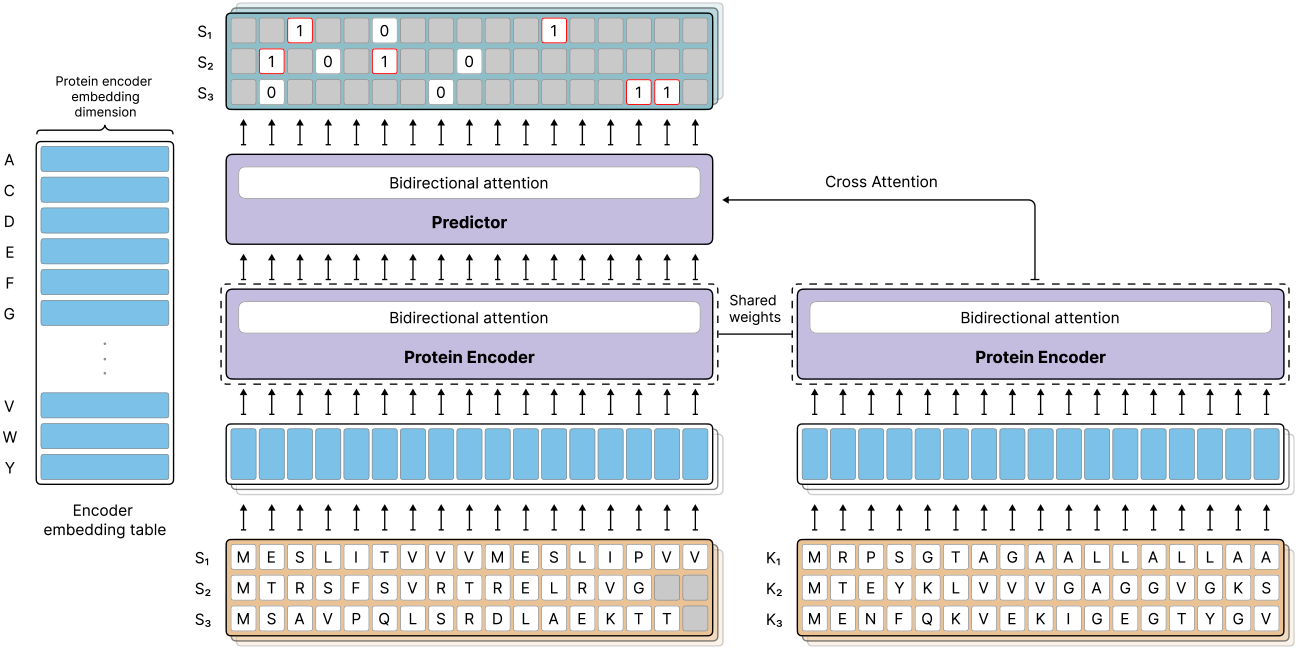
Stage 2 architecture: shared-weight encoders process substrate and kinase sequences; a bidirectional predictor uses cross-attention to output residue-level phosphorylation scores.

The calculated loss for a given sample first uses the standard binary cross-entropy (BCE) formulation. To emphasise hard-to-classify residues while correcting for dataset- and class-specific imbalance, we multiply each residue by a sample weight *w*_*i*_, yielding the weighted loss in (Equation 4). Here *w*_*i*_ ≥ 0 is the product of a dataset-level weight and, when enabled, a positive-class token weight.

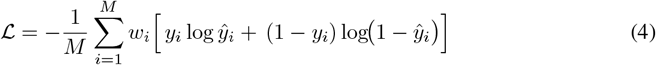

### 2.3 Datasets

We utilized the GPS 6.0 [8] dataset, originally comprising 24 160 p-sites. After preprocessing and mapping IDs to the UniProt database, we retained 13 401 sequences annotated with kinase information, covering 386 kinases across 12 distinct groups validated against Kinase.com for Homo sapiens species. The final dataset contains Uniprotids, substrate sequences, kinase sequences, group information, and p-sites. To reduce sequence similarity, CD-HIT [23] was applied with a 70% threshold. More details of data preparation are described in Appendix A.3.

We prepared an additional test set based on dark kinases provided by [24]. Dark kinases, a subset of human *serine*/*threonine* (`S`/`T`) kinases, remain poorly characterized, with limited knowledge about their substrates, signaling functions, and regulatory mechanisms [25, 24]. To address this gap, over 80 understudied dark kinases were experimentally profiled using positional scanning peptide arrays, which uncovered previously unknown substrate motifs and provided a foundation for functional annotation [24]. For each dark kinase, the highest-scoring position in its position-specific scoring matrix was selected as the predicted p-site. Corresponding substrate sequences and kinase sequences were retrieved from UniProt. The resulting test set contained 8 026 samples representing 29 dark kinases from five distinct kinase groups, with each sample comprising the kinase sequence, its substrate sequence, and the associated p-site.

To evaluate zero-shot performance, we constructed an independent test set comprising dark kinases absent from the GPS dataset (Figure 4c). The set contained 5 391 kinase–substrate pairs, each annotated with the corresponding p-site, and represented 26 distinct kinase types. The frequency of each kinase across the groups, with dark kinases in the training and validation sets highlighted, is shown in Figures 6 and 7.

**Figure 4.**
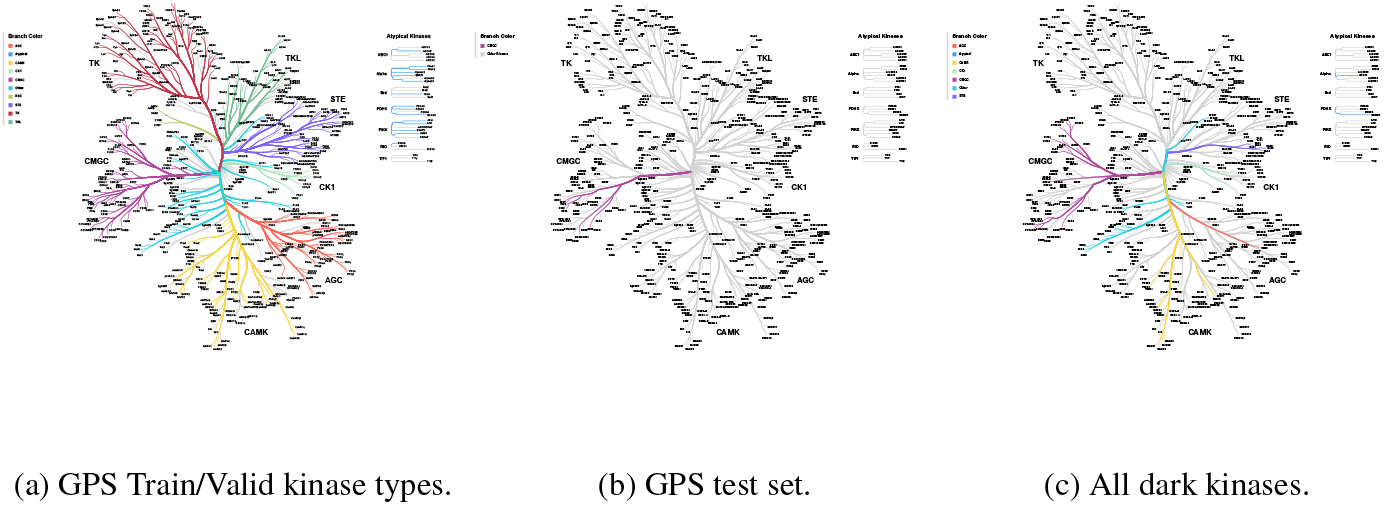
Kinase-type distributions across our datasets, with colors indicating phylogenetic relationships. (a) Distribution of kinase types in the training and validation sets. (b) Kinases belonging to a representative GPS test data. (c) Dark kinases, showing both in-distribution and out-of-distribution cases relative to the training data.

**Figure 5.**
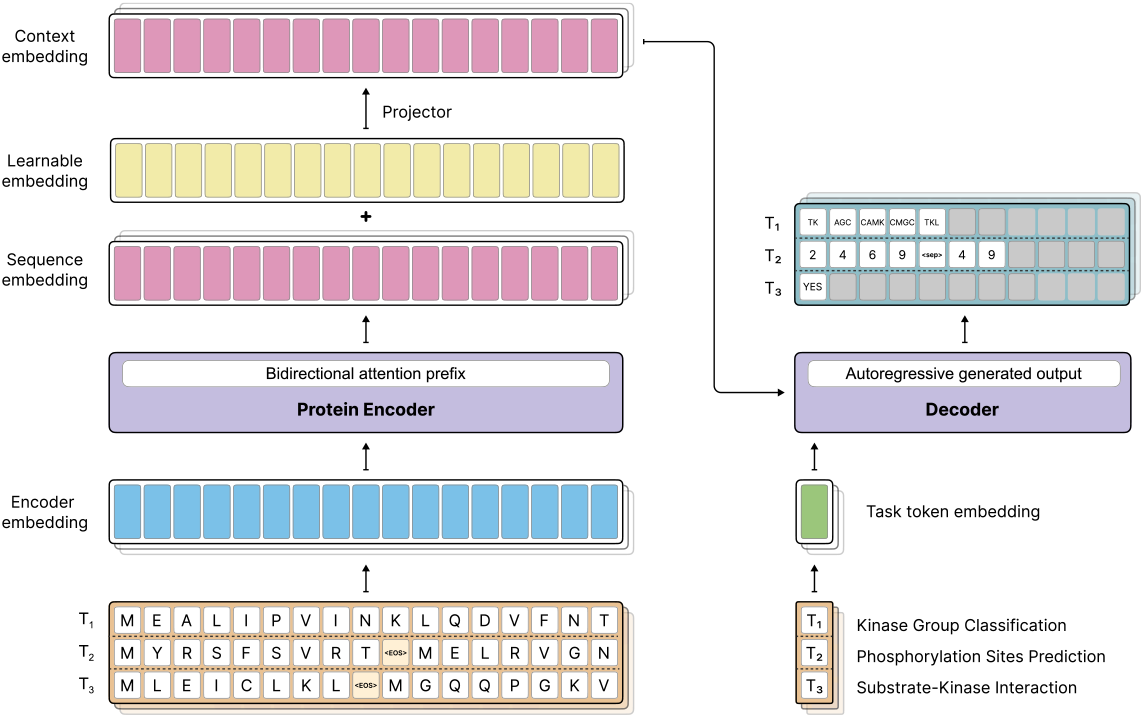
Overview of the proposed model architecture during inference. The bidirectional protein encoder processes substrate and kinase sequences to generate embeddings, which the autoregressive decoder uses to predict output tokens. The figure highlights the integration of the pre-trained ESM-2 model with the autoregressive decoder for kinase group classifications, substrate-kinase interactions, and phosphorylation site prediction.

**Figure 6.**
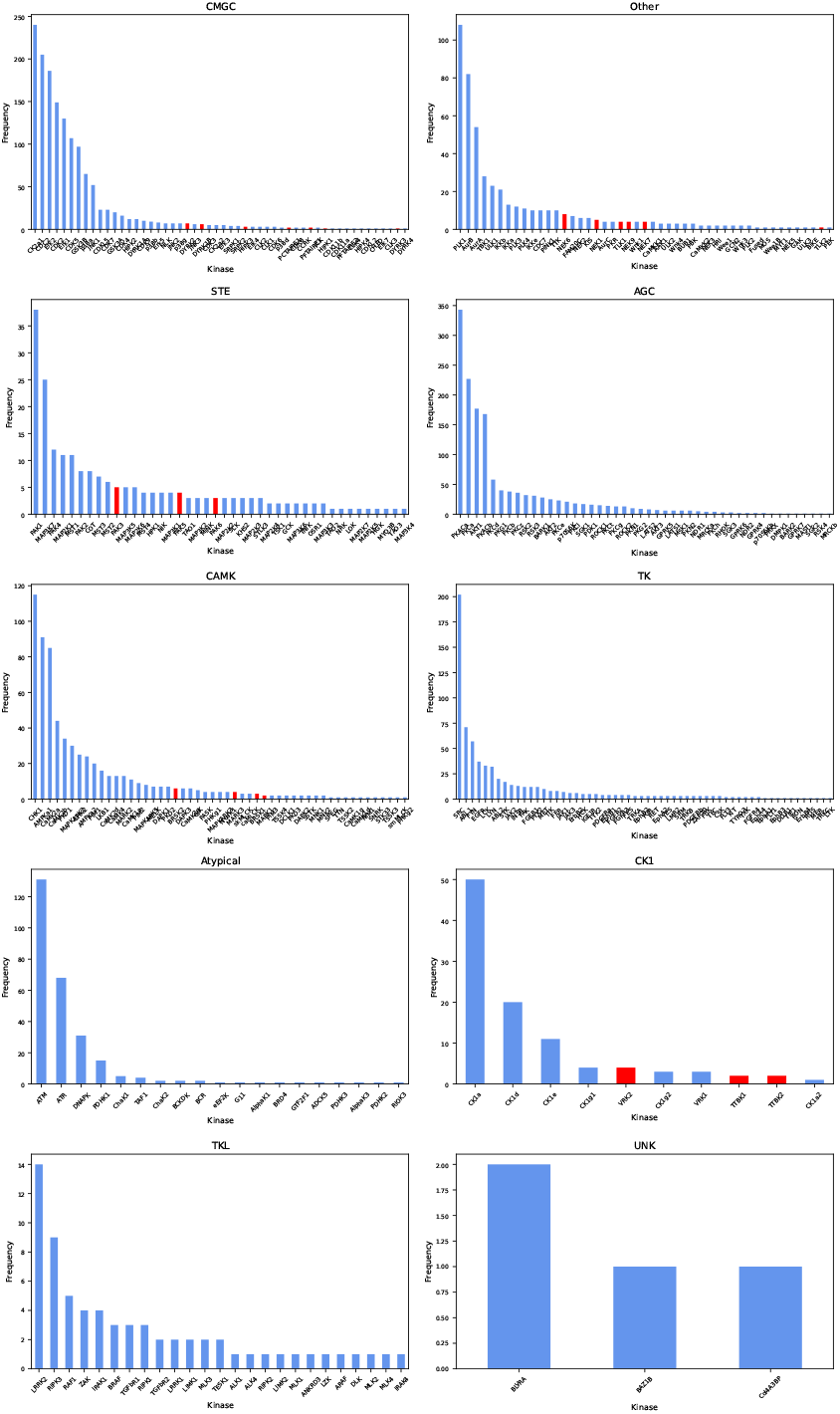
Frequency distribution of kinases across groups in the training dataset, where dark kinases are indicated in red for visual emphasis.

**Figure 7.**
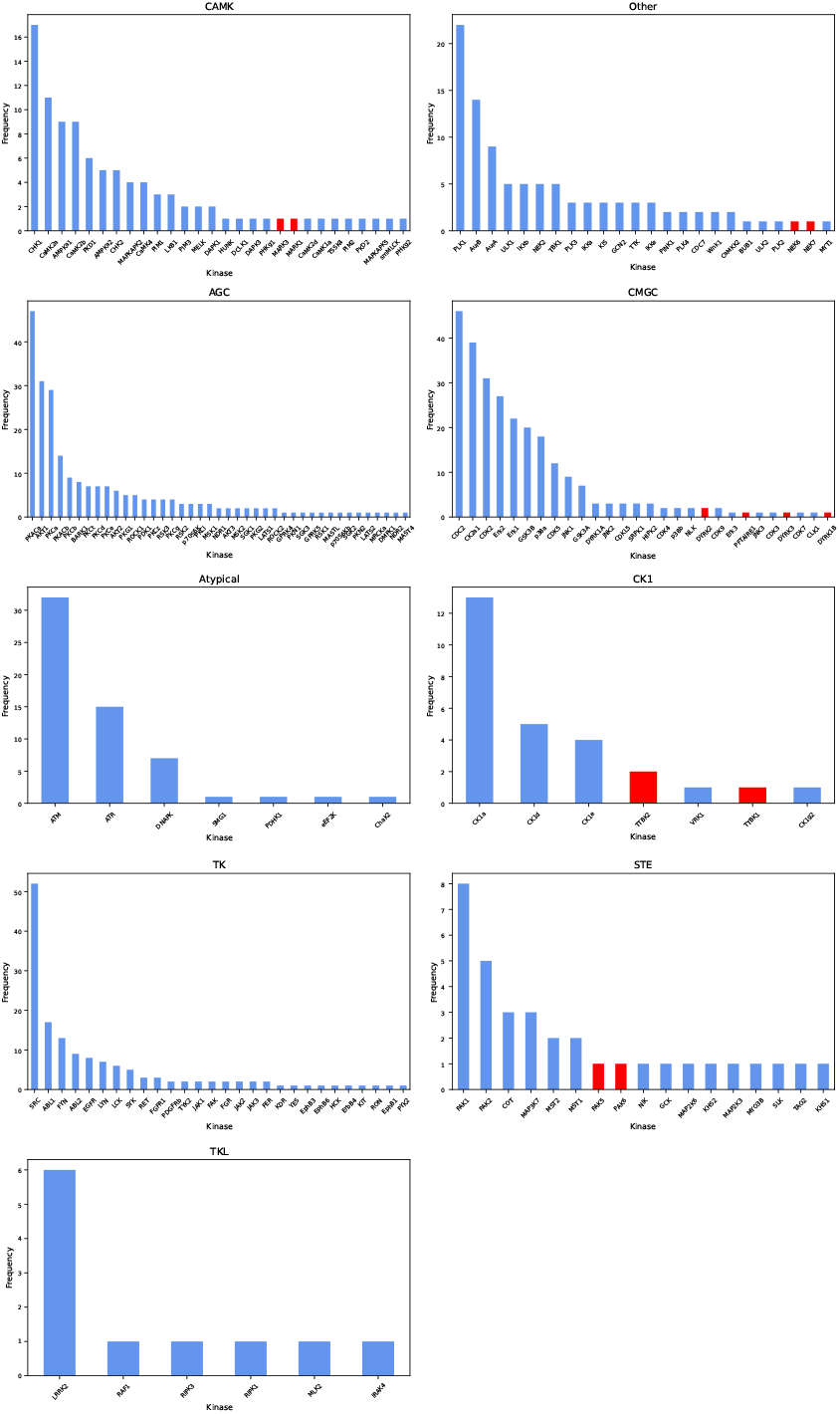
Frequency distribution of kinases across groups in the validation dataset, where dark kinases are indicated in red for visual emphasis.

## 3 Experiments

We structured our experiments to progressively validate the *Stage 1* across the three target tasks. First, we assessed kinase group classification using full substrate sequences alone. Next, we applied hard negative sampling to strengthen the kinase–substrate interaction prediction task. To empower the p-site prediction capability of Prot2Token, we performed self-supervised pre-training of the autoregressive decoder and fine-tuned it for kinase-specific p-site prediction, benchmarking against state-of-the-art methods. Finally, we evaluated the *Stage 2* specialized architecture on the p-site prediction task, including in-distribution and zero-shot dark kinase samples.

For all experiments, we initialized the protein encoder of *Prot2Token* with the pre-trained ESM2 650m model for all three kinase tasks, and the second stage architecture includes ESMC 600m and different scales of ESM-2. We enabled fine-tuning for the weights of the last six blocks of the encoders on all experiments, while keeping the embedding parameters of the encoder frozen unless specified otherwise. All other layers in the model were trained during the experiments. We employed the AdamW optimizer [26] with a weight decay of 0.1, *β*_1_ = 0.9, *β*_2_ = 0.98, and epsilon to 1e-7 as default hyperparameters across all training runs. The learning rate followed a cosine annealing schedule with an initial warm-up phase [27], starting from 1e-6 and increasing to 5e-5 over the first 256 steps unless specified otherwise. For *Stage 2* of training, a weight of 2 was applied to the positive class loss value to handle the label imbalance issue. All training was conducted using the PyTorch 2.6 framework [28] on a single node equipped with 4×A100 80GB Nvidia GPUs.

### 3.1 Kinase Group Classification

In this section, we aimed to predict kinase groups based on substrate sequences. Specifically, we investigated how much information about the related kinase groups the model can infer solely from substrate sequences. To achieve this, we considered our processed training and validation datasets (Appendix A.3), assigning multi-label classification labels by removing *Unknown* group samples from the training set and merging the remaining nine kinase groups associated with each substrate. The model takes a substrate sequence as input and predicts the corresponding kinase groups in alphabetical order. The result is presented in Table 1.

**Table 1:** F1 scores of our method on each kinase group, based on full substrate sequences.

| Group | AGC | Atypical | CMGC | CAMK | CK1 | Other | STE | TK | TKL |
| --- | --- | --- | --- | --- | --- | --- | --- | --- | --- |
| **F1** | 0.5551 | 0.4928 | 0.634 | 0.4196 | 0.5385 | 0.6047 | 0.3571 | 0.6738 | 0.5 |

Next, we compared the embedding representations of all unique kinase sequences before and after fine-tuning the protein encoder part on the kinase group classification model. The fine-tuned model demonstrated slightly better separation of kinase sequences into their respective groups, even though kinase sequences were not explicitly included during training (Table 2). Additional details are provided in Appendix A.4.

**Table 2:**
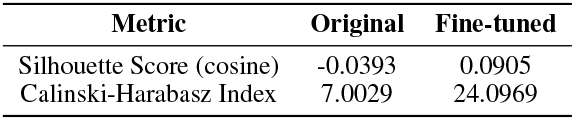
Unsupervised separability metrics of all unique kinase sequences with respect to the original and fine-tuned version of ESM2 650m model.

| Metric | Original | Fine-tuned |
| --- | --- | --- |
| Silhouette Score (cosine) | -0.0393 | 0.0905 |
| Calinski-Harabasz Index | 7.0029 | 24.0969 |

### 3.2 Kinase-Substrate Interaction Prediction

Although predicting protein–kinase interactions is a meaningful task, its practical utility is somewhat diminished, as such interactions can often be inferred implicitly through p-site prediction. Nevertheless, we include this task for completeness and comparative evaluation. We formulated the problem as determining whether a given protein substrate interacts with a specific kinase sequence. To construct training data, we created both positive and hard negative samples from the dataset. However, generating biologically plausible negative labels involves complex and labor-intensive procedures, which are noisy and not easily scalable. As such, we have relegated the details of negative sample construction to Appendix A.4. The resulting performance is reported in Table 3.

**Table 3:** The details of protein-kinase interaction performance versus other methods on the positive and randomly selected negative samples from the validation set.

| Metric | Ours | GPS 6.0 [8] | PhosphormerST [29] |
| --- | --- | --- | --- |
| Accuracy (%) | **72** | 58 | 49 |
| F1 | **0.71** | 0.61 | 0.603 |

### 3.3 Kinase-Substrate Phosphorylation Site Prediction

In this section, to adapt Prot2Token imitate p-site prediction, a self-supervised pre-training stage was incorporated. We randomly sampled 4 million protein sequences from the UniRef50 database for training and 4 000 sequences for validation. From these, we artificially generated 80 million training samples and 20 000 validation samples by treating each amino acid type within a protein as an individual sample. Subsequently, we further randomly sampled 1 million training and 1 000 validation samples to construct the final datasets of this part. For training, we used an input sequence length of 1 280, a weight decay of 0.01, and a batch size of 192 samples, equivalent to 73 728 tokens. The warm-up phase consisted of 512 steps. During training, we froze all the encoder weights while allowing all other parameters to be updated. After 16 epochs, the model achieved a validation perplexity of 2.31, indicating that it could almost perfectly reconstruct protein sequences from the encoder’s embeddings.

Building on our ability to predict protein-kinase interactions, we extended our approach to precise phosphorylation site prediction. To achieve this, we selected all protein-kinase sequence pairs along with their corresponding phosphorylation sites and jointly trained them alongside 20 self-supervised tasks. This fine-tuning phase utilized the latest checkpoint from the self-supervised pre-training stage as its initial checkpoint. For this phase, we reduced the number of self-supervised tasks to a total of 20 000 samples. Additionally, substrate sequences exceeding 1 280 amino acids in length were excluded during training and evaluation. The results are shown in Table 4. More details are presented in A.4. To further enhance phosphorylation site prediction, we trained the *Stage 2* architecture using the same kinase–substrate pairs and phosphorylation site annotations, and demonstrated improved performance compared to *Stage 1*. The result of the *Stage 2* model is presented in Table 4. Additionally, we conducted ablation studies on the *Stage 2* model to assess the impact of different protein language model backbones. The results, summarized in Table 5.

**Table 4:** Comparative results of our method against leading tools for p-sites prediction (KinasePhos3, NetPhose 3.1, MusiteDeep and GPS 6.0) across the validation and GPS test. NetPhose 3.1 and Musite did not support all kinase groups in the validation set. *†*: the decoder initialized from scratch.

| Method | Validation Set |  |  | GPS Test Set |  |  |
| --- | --- | --- | --- | --- | --- | --- |
|  | Precision | Recall | F1 | Precision | Recall | F1 |
| KinasePhos3 [9] | 0.0773 | **0.9557** | 0.138 | 0.0215 | **0.9856** | 0.0421 |
| NetPhos 3.1 [22] | - | - | - | 0.0325 | 0.0137 | 0.0193 |
| GPS 6.0 [8] | 0.2323 | 0.4549 | 0.3076 | 0.1564 | 0.5054 | 0.2398 |
| Musite [11] | - | - | - | 0.1348 | 0.8057 | 0.2310 |
| Stage 1 (Ours)† | 0.0403 | 0.0016 | 0.0206 | 0.0131 | 0.0033 | 0.007 |
| Stage 1 (Ours) | **0.5753** | 0.4411 | 0.4994 | 0.4575 | 0.3660 | 0.4069 |
| Stage 2 (Ours) | 0.4757 | 0.4637 | **0.4696** | **0.4597** | 0.4302 | **0.4444** |

**Table 5:** Effect of encoder scales of Stage 2 architecture on kinase-substrate phosphorylation task.

| PLM Backbone | F1 |
| --- | --- |
| ESM-2 35m | 0.3211 |
| ESM-2 150m | 0.3806 |
| ESM-2 650m | 0.4260 |
| ESMC 600m | 0.4444 |

### 3.4 Prediction of Dark Kinases

To evaluate the dark kinases included in our dataset and enable a direct comparison with GPS 6.0, we performed inference using our best model on these kinases and compared the results with other p-site prediction methods. All 29 selected dark kinases were also supported by GPS 6.0, ensuring a fair comparison. Predictions from GPS were obtained using medium thresholds, and the results are summarized in Table 6a.

**Table 6:**
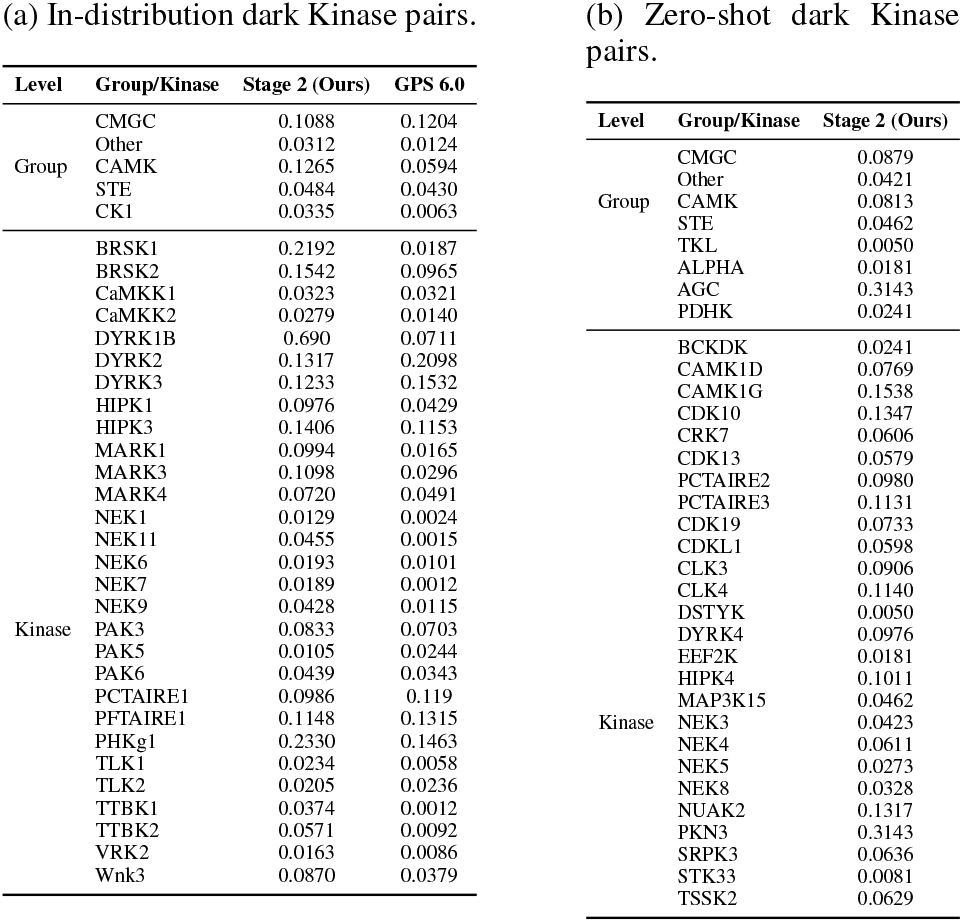
Dark kinase in-distribution (a) and out-distribution (b) evaluation performance of kinase-substrate p-site prediction, based on F1-score metric.

**Table 7:** Key hyperparameters for ProtToken architecture.

|  |  |
| --- | --- |
| **Protein encoder** | ESM-2 650m |
| **Protein encoder dimension** | 1 280 |
| **Fine-tuned blocks** | 6 |
| **Decoder dimension** | 640 |
| **Decoder block** | 16 |
| **Decoder heads** | 16 |
| **Decoder FF dimension** | 2560 |
| **Decoder activation** | GELU |

**Table 8:** Key hyperparameters for Stage 2 architecture across different model scales.

|  |  |  |  |  |
| --- | --- | --- | --- | --- |
| **Protein encoder** | ESM-2 35m | ESM-2 150m | ESM-2 650m | ESMC 600m |
| **Protein encoder dimension** | 480 | 640 | 1 280 | 1 152 |
| **Fine-tuned blocks** |  |  | 6 |  |
| **Decoder dimension** | 480 | 640 | 1 280 | 1 152 |
| **Decoder block** |  |  | 6 |  |
| **Decoder heads** |  |  | 8 |  |
| **Decoder FF dimension** |  |  | 640 |  |
| **Decoder activation** |  |  | GELU |  |

While tools like GPS offer limited coverage for dark kinases, our model demonstrates better capabilities even for poorly characterized enzymes. To evaluate this, we performed zero-shot prediction on 26 dark kinases not seen during training. Results are presented in table 6b.

## 4 Discussion

Our results show that substrate sequences alone encode discriminative signals for kinase-group assignment, even in the absence of kinase sequences. This finding implies that substrates carry motif- and context-level cues sufficient to prioritize a small set of candidate kinase families for follow-up, potentially reducing the number of targeted assays needed to confirm phosphorylation events.

Augmenting substrates with kinase sequences, we introduced two complementary architectures. *Stage 1* extends *Prot2Token* with a decoder pre-training task that learns position-aware residue representations from encoder features, yielding substantial gains for kinase-specific site prediction. *Stage 2* replaces causal decoding with bidirectional cross-attention over encoder features, further improving precision–recall trade-offs and delivering state-of-the-art performance on standard benchmarks.

A key outcome is the sign of generalization beyond well-studied kinases. The specialized model (*Stage 2*) retains better performance on understudied dark kinases and, in zero-shot evaluations on unseen kinase–substrate pairs, exhibits the first evidence of zero-shot kinase-specific phosphorylation-site prediction, to our knowledge. Given the central role of dysregulated kinase signaling in oncogenesis and therapy resistance, these capabilities may accelerate hypothesis generation for pathway mapping, biomarker nomination, and kinase prioritization in cancer settings.

## A Appendix

### A.1 Related work

We have categorized approaches in this domain into two main areas: those focused on kinase-specific phosphorylation site prediction and those centered on kinase-protein interaction prediction.

#### A.1.1 Kinase-Specific Phosphorylation Sites

In the first category, studies focus on predicting phosphorylation sites specific to particular kinases, kinase families or kinase groups. These methods often leverage sequence-based features, structural properties, and evolutionary information to identify phosphorylation sites associated with a specific kinase. In this category family of algorithms, GPS [31–35, 8] uses several models integrated to predict different groups and number of p-sites for each peptide with size of 15 amino acids. KinasePhos3.0 [9] used 771 predictive models, developed at different levels, including kinase group, family, and individual kinase levels (SVM and XGBoost) with SHAP-based features to predict kinase-specific phosphorylation sites from 15-residue sequence windows surrounding phosphorylation sites, to predict number of positions and different kinase levels, MusiteDeep [11] used convolutional neural network, DeepPhos [36] used densely connected convolutional neural network with different window sizes of 25, 33, 51 peptides, PhosIDN [18] integrates sequence features extracted through a self-attention-enhanced CNN and protein-protein interaction embeddings processed via a deep neural network, combining them with a bilinear module to predict general and kinase-specific phosphorylation sites.

##### A.1.2 Kinase-Substrate Interaction

The second category addresses the broader challenge of protein-kinase interaction prediction, where the goal is to identify interactions between kinases and their substrate proteins. This task often involves integrating sequence embeddings, structural data, and sometimes even contextual biological information to predict interaction patterns. Phosphormer-ST [29] and Phosphormer [37] both tried to predict substrate-kinase interaction. Phosphormer used transformer-based architecture and Phosphormer-ST finetuned on ESM2-650M [38] parameters. It uses a shared encoder to generate embeddings for both kinase and peptide sequences.

#### A.2 Architecture

#### A.3 Datasets

The whole dataset was gathered from GPS, where its information on kinase families and groups is based on the kinome of *Homo sapiens*. All kinase sequences were extracted from Uniprot DB [39] and Kinase.com. To create a diverse and non-redundant dataset, we applied CD-HIT clustering with a 70% sequence similarity threshold, grouping similar protein substrate sequences and retaining representative sequences from each cluster by removing similar substrate sequences. Representative positive pairs were chosen using the following criteria:

- **Cross-cluster selection:** Substrate-kinase pairs spanning different clusters were retained to preserve diversity across the dataset.
- **Within-cluster selection:** For substrates within the same cluster, only one unique kinase pair was kept, minimizing redundancy while ensuring distinct associations.

Each substrate is treated as an individual sample for model input, with one or more group labels and corresponding phosphorylation sites assigned per sample. Since substrates can be associated with multiple kinase groups and contain multiple phosphorylation sites, the problem is naturally formulated as a multilabel classification task. After preprocessing, the final dataset consisted of 5 385 unique substrates for training and 969 unique substrates for validation. To ensure rigorous evaluation, we defined three distinct test sets, carefully designed to prevent any data contamination between the test, training, and validation sets:

##### Rare-Group

This set includes 14 samples from two rare kinase groups, *RGC* and *PKL*, which have a limited number of available samples. These groups were completely excluded from the training, validation, and test sets.

##### GPS-Test Set

To have a direct comparison with existing methods such as GPS 6.0, we adopted the test set used in the GPS study. This dataset contains 146 samples of substrate-kinase pairs, including phosphorylation site (p-site) and kinase group annotations. All samples belong to the *CMGC* kinase group.

##### Validation Set

This set was created using a random split strategy, ensuring a balanced distribution of kinase groups across both the test and training sets. Additionally, substrates in this set were selected to have minimal sequence similarity to each other, providing a robust measure of the model’s generalization performance. Table 9 presents the number of samples in each set, while Table 10 details the distribution of samples across kinase groups in each dataset.

**Table 9:** Dataset statistics, including the number of samples, phosphorylation sites (p-sites), and kinase groups for the training, validation, GPS test, and rare group test sets, along with overall dataset totals.

| Dataset | Number of samples | Number of p-sites | Number of groups |
| --- | --- | --- | --- |
| All samples | 6 514 | 13 401 | 12 |
| Training set | 5 385 | 10 621 | 10 |
| Validation set | 969 | 2 455 | 9 |
| GPS-test | 146 | 300 | 1 |
| Rare-Group | 14 | 25 | 2 |

**Table 10:** Distribution of samples across kinase groups for the training, validation, GPS test, and rare group test sets.

| Group | Training set | Validation set | GPS-test | Rare-Group |
| --- | --- | --- | --- | --- |
| AGC | 1 446 | 231 | - | - |
| Atypical | 270 | 58 | - | - |
| CAMK | 653 | 96 | - | - |
| CK1 | 100 | 27 | - | - |
| CMGC | 1 466 | 264 | 146 | - |
| Other | 491 | 99 | - | - |
| STE | 211 | 34 | - | - |
| TK | 677 | 149 | - | - |
| TKL | 68 | 111 | - | - |
| RGC | - | - | - | 2 |
| PKL | - | - | - | 12 |
| UNK | 4 | - | - | - |

While *serine* (`S`) and *threonine* (`T`) are the primary residues studied in phosphorylation research and commonly targeted by prediction tools, other amino acids—such as *histidine* (`H`) and *aspartate* (`D`)—are also known to undergo phosphorylation, particularly in prokaryotic systems and specific eukaryotic contexts. In our datasets, we identified phosphorylation sites on additional amino acids. We excluded these sites from our positive sites and finalize each set. Table 11 presents the distribution of phosphorylation sites exclusively on `S` and `T` residues.

**Table 11:** Distribution of positive sites across kinase groups for the training, validation, GPS test, and rare group test sets.

| Dataset | Serine (S) | Threonine (T) |
| --- | --- | --- |
| Training set | 5 726 | 1 874 |
| Validation set | 1 472 | 465 |
| GPS Test | 201 | 76 |

#### A.4 Experiments

##### A.4.1 Kinase Groups Classification

We analyzed the sequence embeddings of unique kinase sequences extracted from all samples in the GPS 6.0 dataset. For this analysis, kinase sequences from the *RGC, PKL*, and *UNK* groups were excluded. These sequences were processed using the pre-trained ESM-2 650M model to generate token-wise embeddings, with a maximum sequence length of 2 048. After extracting the model’s outputs, we removed the beginning-of-sequence (BOS) and end-of-sequence (EOS) tokens and applied average pooling to obtain fixed-length representations of dimension 1 280, matching the model’s embedding size.

Next, we employed t-SNE and UMAP for dimensionality reduction, enabling visualization of the embeddings in a two-dimensional space according to their group assignments. Given that the group labels for the kinase sequences were known, these labels were used to show the clusters visually on t-SNE and UMAP graphs with different colors. Also, we calculated unsupervised clustering metrics, such as the silhouette score and the Calinski-Harabasz index. We repeated the entire process for the fine-tuned ESM-650M checkpoint that is trained on the training set of kinase group classification labels. Similar to the first part, we performed dimensionality reduction visualizations and computed clustering metrics to evaluate the differences of both visual and unsupervised clustering metrics. The results are illustrated in Figures 9 and 8 and Table 2.

**Figure 8.**
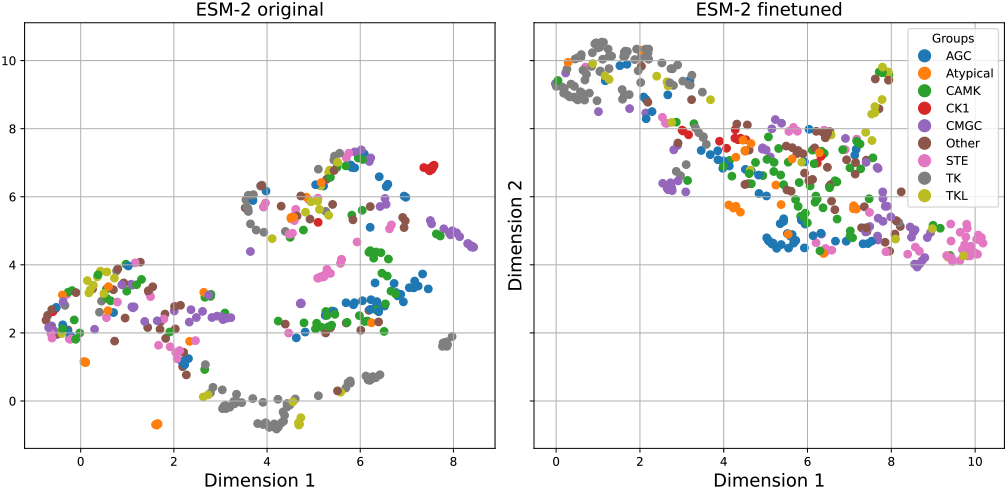
UMAP visualization of unique kinase sequences on the original and fine-tuned checkpoints of ESM-2 650m

**Figure 9.**
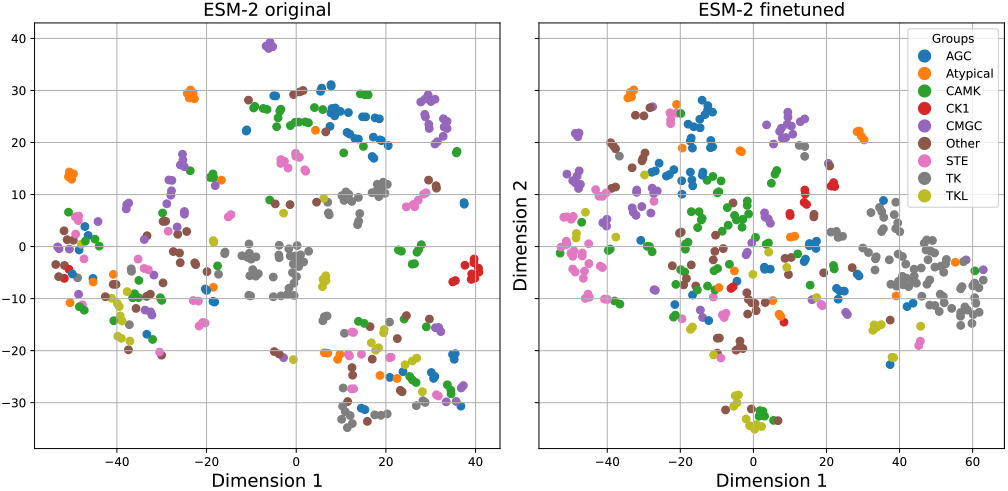
t-SNE visualization of unique kinase sequences on the original and fine-tuned checkpoints of ESM-2 650m.

##### A.4.2 Protein-Kinase Interaction

The positive data pairs are unique protein-kinase pairs with information of full protein sequence and full kinase sequence. The novelty of our method lies in the strategic selection of hard negative samples. Inspired from data pruning [15] method, hard negative samples preparation relies on choosing kinase sequences with embeddings that exhibit minimal Euclidean distances to other kinases.

To construct the negative pairs, we iterated through all unique substrates in the dataset, processing each substrate individually. For each substrate, we first identified its associated positive kinases from the positive pairs. Using pre-computed embeddings, we calculated the Euclidean distance between each unique kinase embedding in the dataset to form the distance map. From these distances, we selected the top *k* closest kinases, excluding the positive kinase itself, to form *k* hard negative pairs for the substrate (Figure 10). This process was repeated for every substrate, ensuring that each one was paired with both its positive kinases and the most challenging negative kinases based on all kinase embeddings. By focusing on embedding distances to identify closely related negatives, this approach ensured a challenging dataset that effectively trained the model to distinguish between positive and negative pairs. We evaluated the impact of training protein-kinase interaction with hard negative samples versus random negative samples, focusing exclusively on the positive samples in the validation set. The results are summarized in Table 12.

**Table 12:** Comparison of different negative sampling strategies for protein-ligand interaction training, evaluated on the positive samples of the validation set.

| Negative sample strategy | Random | Hard |
| --- | --- | --- |
| **F1 score** | 0.771 | 0.8325 |

**Figure 10.**
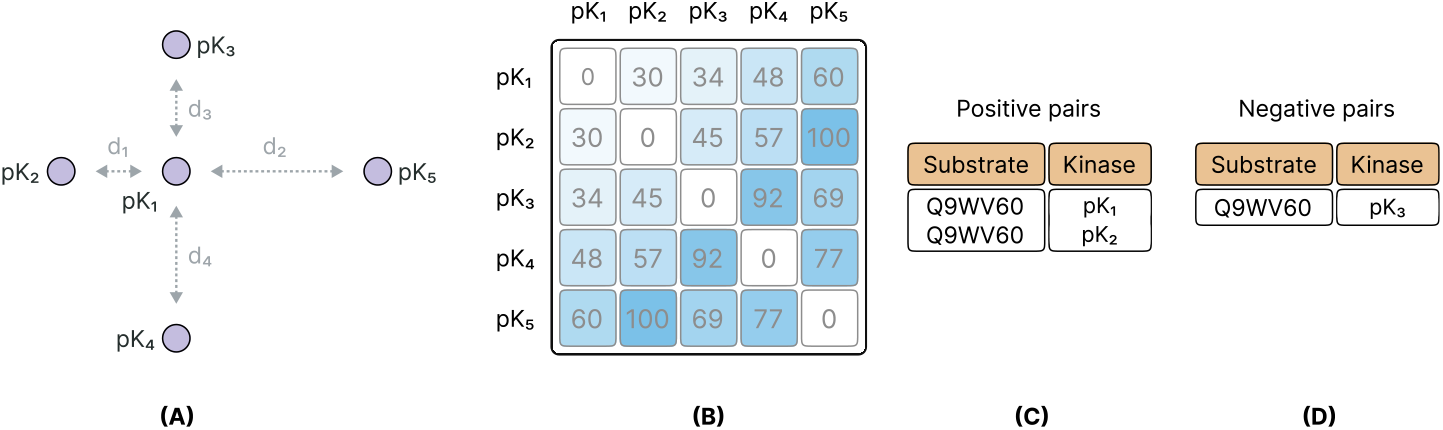
(A) Visualization of kinase embedding distances: The diagram depicts a substrate (center, *pK*_1_ and its surrounding kinases (*pK*_2_, *pK*_3_, *pK*_4_), and *pK*_5_, illustrating the Euclidean distances (*d*_1_, *d*_2_, *d*_3_, *d*_4_) between their embeddings in the learned space. (B) Distance matrix of kinase embeddings: A heatmap representing pairwise Euclidean distances between kinase embeddings. Lighter shades correspond to shorter distances, indicating higher similarity, while darker shades represent greater dissimilarity. (C) Positive sample pairs: Examples of positive protein-kinase pairs (e.g., substrate Q9WV60 with kinases *pk*_1_ and *pk*_2_, representing known functional phosphorylation interactions. (D) Negative sample pairs: Examples of hard negative protein-kinase pairs (e.g., substrate Q9WV60 with kinase *pk*_3_, selected based on minimal Euclidean distances in the embedding space to encourage robust discriminative learning.

During the training, we set a maximum context length of 1 280 tokens for the combined substrate-kinase sequences, truncating them when necessary to fit within this limit.

We evaluated GPS 6.0 [8] and PhosphormerST [29] using our own validation set. For GPS 6.0, substrate sequences were provided as input, and in contrast to PhosphormerST, GPS 6.0 does not support kinase domain sequence as the input. To add the information of kinases during the prediction of GPS 6.0, we have selected each group as the kinase information in their webserver. The method generated a score and cutoff value for each phosphorylation site prediction. An interaction between the substrate and kinase was valid if the model predicted at least one phosphorylation site in the substrate sequence. On the other hand, PhosphormerST takes peptides and kinase domains as inputs and generates a prediction score for each sample. We created peptides with a length of 15 and mincluded kinase domain information as the other part of the input. Interactions were considered valid if the score exceeded 0.5.

#### A.5 Kinase-Substrate Phosphorylation Site Prediction

During the *Stage 1* experiment, we set the total sequence length (including both substrate and kinase sequences) to 2 048 tokens, truncating kinase sequences as needed to fit within this limit. The batch size was configured to accommodate 98 304 tokens per iteration. It is important to highlight that, although the self-supervised tasks could have been entirely excluded from the fine-tuning stage, retaining a subset of these samples led to a noticeable improvement in the model’s performance on protein-kinase phosphorylation site prediction. We also found that without the self-supervised checkpoint, the model’s performance in phosphorylation site prediction dropped sharply, reaching an F1 score of less than 0.1 on the validation set, highlighting the necessity of pre-training for maintaining predictive accuracy. The max input size of the *Stage 2* architecture was set to 1 280 tokens.

We compared our results with two phosphorylation prediction tools, GPS 6.0 and KinasePhos3 [9]. We used Medium threshold for GPS and scores more than 0.5 for all other tool, and the predicted phosphorylation sites were compared to experimentally validated sites. To generate the results, we selected each kinase group individually on the tools. However, there is a strong possibility of *data contamination* between our validation set and the GPS 6.0 training set. As a result, GPS 6.0 may achieve artificially high performance on our validation set due to memorization, while its real-world performance on unseen substrates could be even lower.

#### A.6 Availability of Resources

To support further research and development in the field, we make our codes, trained models, datasets, and a Python package publicly available for use by researchers and the broader scientific community. These resources can be accessed at the related GitHub repository.

## Footnotes

‡ In this paper, we refer to it as kinase-substrate phosphorylation.

